# Perceptual consequences of interocular imbalances in the duration of temporal integration

**DOI:** 10.1101/2022.02.16.480712

**Authors:** Benjamin M. Chin, Johannes Burge

## Abstract

Temporal differences in visual information processing between the eyes can cause dramatic misperceptions of motion and depth. Processing delays between the eyes cause the Pulfrich effect: oscillating targets in the frontal plane are misperceived as moving along near-elliptical motion trajectories in depth (Pulfrich, 1922). Here, we explain a previously reported but poorly understood variant: the anomalous Pulfrich effect. When this variant is perceived, the illusory motion trajectory appears oriented left- or right-side back in depth, rather than aligned with the true direction of motion. Our data indicate that this perceived misalignment is due to interocular differences in neural temporal integration periods, as opposed to interocular differences in delay. For oscillating motion, differences in the duration of temporal integration dampen the effective motion amplitude in one eye relative to the other. In a dynamic analog of the Geometric effect in stereo-surface-orientation perception (Ogle, 1950), the different motion amplitudes cause the perceived misorientation of the motion trajectories. Forced-choice psychophysical experiments, conducted with either different spatial frequencies and/or different onscreen motion damping in the two eyes, show that the perceived misorientation in depth is associated with the eye having greater motion damping. A target-tracking experiment provided more direct evidence that the anomalous Pulfrich effect is caused by interocular differences in temporal integration and delay. These findings highlight the computational hurdles posed to the visual system by temporal differences in sensory processing. Future work will explore how the visual system overcomes these challenges to achieve accurate perception.

## Introduction

Temporal processing changes with the sensory stimuli being processed. Some sensory signals take longer to process than others. Stimulus-based differences in temporal processing delays— relative latencies—have received significant attention in vision science and neuroscience. Luminance signals are processed more quickly than chromatic signals. High luminance signals are processed more quickly than low luminance signals. High contrast signals are processed more quickly than low contrast signals. And low frequency stimuli are processed more quickly than high frequency stimuli. Despite differences in the speed by which these signals are processed, they are integrated by the brain. The computational rules that govern the integration of complementary signals with different temporal dynamics are not yet well understood. Identifying striking perceptual phenomena that result from combining such signals, and developing high-fidelity tools for measuring and characterizing these phenomena, should aid the discovery of computational principles underlying the combination rules.

Binocular integration of information between the eyes is crucial to depth perception. When a scene is viewed binocularly, the images are different in the two eyes because of their different vantage points on the scene. The spatial differences between the images in the two eyes underlie stereopsis, the perception of depth from binocular information. Estimation of these spatial differences can be impacted by differences in temporal processing between the eyes, especially when the images move.

Simple processing delays between the eyes cause oscillating targets in the frontal plane to be misperceived as moving along near-elliptical motion trajectories in depth (Fig. 1AB). Such interocular delays cause effective spatial displacements in one eye relative to the other—a neural disparity—that results in the illusory motion in depth. This illusion is known as the Pulfrich effect (Pulfrich, 1922). Luminance, contrast, and blur differences between the eyes are all known to the effect. Two types of the Pulfrich effect have been reported: the classic Pulfrich effect and the reverse Pulfrich effect. In the classic Pulfrich effect, the eye with lower luminance or contrast is processed more slowly (Lit, 1949; Reynaud & Hess, 2017; Wilson & Anstis, 1969; Fig. 1A). In the reverse Pulfrich effect, the eye with lower image quality (due to blur) is processed more quickly (Burge, Rodriguez-Lopez, & Dorronsoro, 2019; Rodriguez-Lopez, Dorronsoro, & Burge, 2020; Fig. 1B).

**Figure 1.**
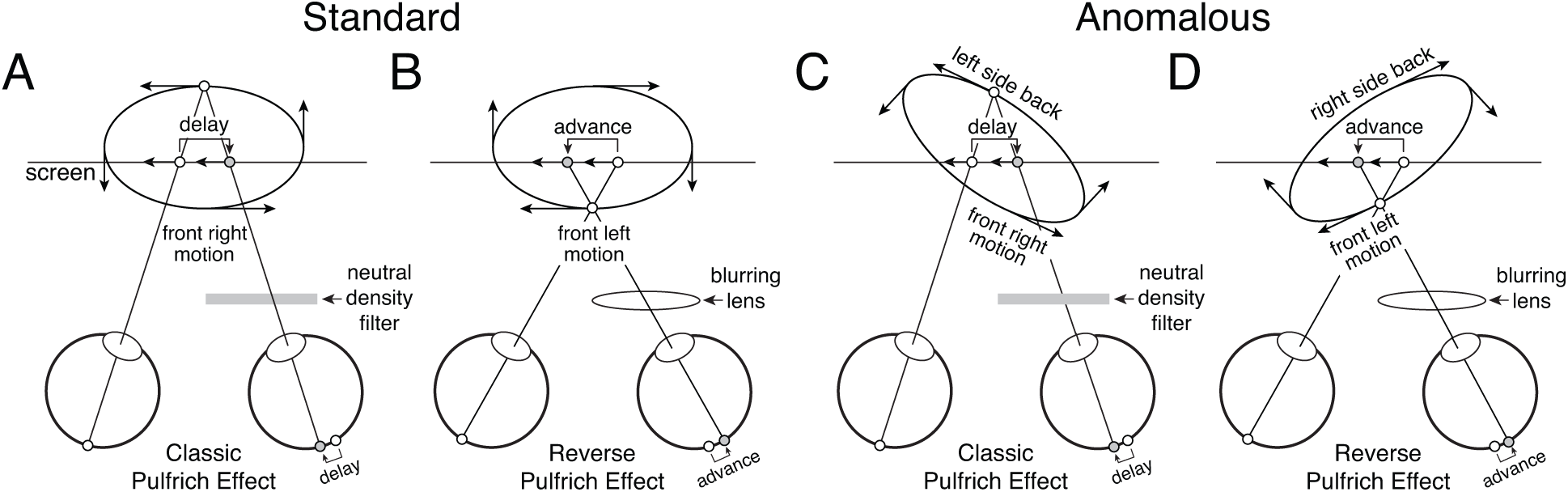
Standard and anomalous versions of the classic and reverse Pulfrich effects. **A** Standard version of the classic Pulfrich effect. A neutral density filter delays the signal in one eye relative to the other by decreasing luminance. **B** Standard version of the reverse Pulfrich effect. A blurring lens advances the signal in one eye relative to the other. **C** Anomalous version of the classic Pulfrich effect. **D** Anomalous version of the reverse Pulfrich effect.

The reverse Pulfrich effect is mediated by blur-induced differences in the spatial frequency content between the stimuli in the two eyes (Burge et al., 2019; Rodriguez-Lopez et al., 2020). Blurring an image low-pass filters the image: high spatial frequencies are selectively removed. Because high spatial frequencies are processed more slowly than low spatial frequencies, the sharp image is processed more slowly than the blurry image. Complementarily, high-pass filtering increases the proportion of high frequencies in the image, and causes the high-pass filtered image to be processed more slowly (Burge et al., 2019). Similarly, if the two eyes are stimulated by moving Gabor stimuli with different carrier frequencies, signals from the eye with higher frequencies are processed more slowly (Min, Reynaud, & Hess, 2020). Thus, simple processing delays (i.e. time shifts in neural responses) nicely account for the standard Pulfrich effect: the perception of illusory 3D motion aligned with the true path of motion.

Anomalous Pulfrich percepts have also been reported (Emerson & Pesta, 1992; Harker & O’neal, 1967; Trincker, 1953; Weale, 1954). In such cases, observers report perceiving near-elliptical motion paths with principal axes that are rotated in depth relative to the true direction of motion (Fig. 1CD). Simple processing delays cannot explain these percepts. Various explanations have been proposed regarding the cause of anomalous Pulfrich percepts: saccadic suppression, velocity extrapolation, and perceptual distortion of objective visual space (Emerson & Pesta, 1992; Harker & O’neal, 1967; Trincker, 1953; Weale, 1954). But scientific consensus has not coalesced around any of these explanations. The aim of this paper is to explain this previously reported but poorly understood variant of the illusion.

We hypothesize that anomalous Pulfrich percepts—illusory motion trajectories that are rotated in depth with respect to the true motion trajectory—are caused by differences in the duration of time over which each eye integrates visual information; that is, different temporal integration periods. To understand this hypothesis, consider the temporal dynamics of sensory processing. The neural response to a sensory stimulus evolves over time. This temporal evolution can be described by an impulse response function. For a moving stimulus, the effective position over time of the neural image is impacted by this impulse response function. If the impulse response function in one eye is delayed relative to the impulse response function in the other eye (i.e. they are time-shifted copies of each other), stereo-geometry predicts the standard Pulfrich effect, when oscillatory motion is presented (Fig. 2AB). If, on the other hand, the impulse response function in one eye is both delayed and has a longer temporal integration period than the impulse response function in the other eye, then the amplitude of the effective motion signal will be damped in that eye relative to the other eye (Fig. 2CD). In this case, stereo-geometry predicts the anomalous Pulfrich effect: illusory motion-in-depth along a trajectory that is misaligned with the true direction of motion (see Fig. 1CD and Discussion).

**Figure 2.**
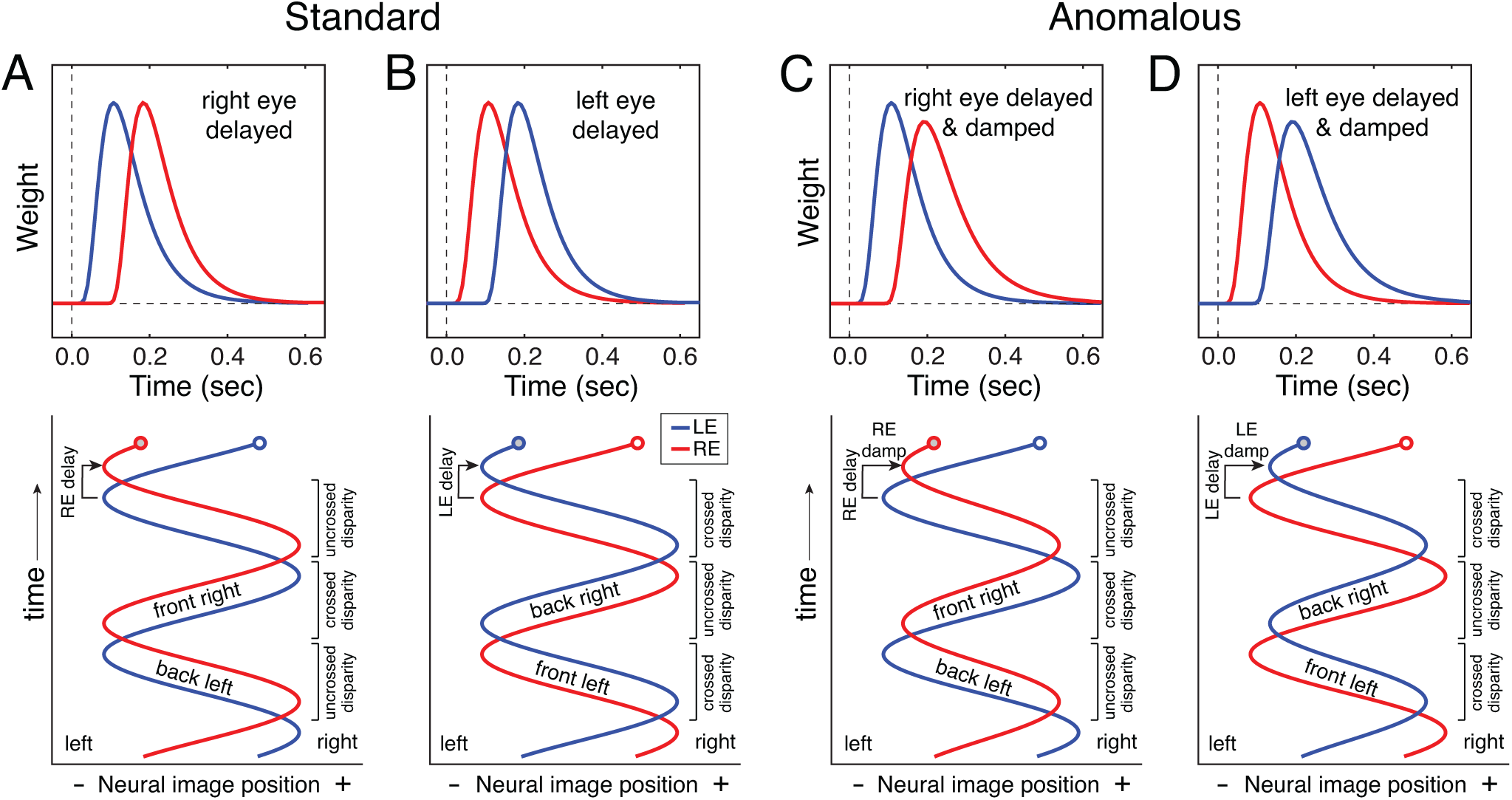
Predicting standard and anomalous Pulfrich percepts. Temporal impulse response functions (top) and effective neural image positions over time (bottom) for the left (blue) and right (red) eyes when **A** processing in the right eye is delayed relative to the left, **B** processing in the left eye is delayed relative to the right, **C** processing in the right eye is delayed and damped (due to a longer temporal integration period) relative to the left, and **D** processing in the left eye is delayed and damped relative to the right. Standard Pulfrich percepts result from delays only. Anomalous Pulfrich percepts result when the effective motion trajectory in one eye is both delayed and damped relative to the other (see Discussion).

Informally, we have most often observed anomalous Pulfrich percepts when there is different spatial frequency content in the two eyes. It is well-known that different spatial frequencies are processed both with different delays and with temporal integration periods of different durations. Neurons in early visual cortex (V1) and the middle-temporal area (MT) respond to higher spatial frequencies with more delay and longer temporal integration periods, all else equal (Bair & Movshon, 2004; Frazor, Albrecht, Geisler, & Crane, 2004; Vassilev, Mihaylova, & Bonnet, 2002). Psychophysical experiments have shown that human perceptual responses are similarly affected by spatial frequency (Levi, Harwerth, & Manny, 1979).

Here, with a traditional two-alternative forced choice (2AFC) paradigm previously used to study the Pulfrich effect, we first presented different spatial frequencies to the two eyes and asked observers to report the perceived orientation of the motion trajectory in depth (‘left side back’vs ‘right-side back’; see Fig. 1CD). Anomalous (i.e. non-fronto-parallel) motion trajectories were reported in the expected direction. Next, to confirm that effective motion damping in the eye with the higher spatial frequency was indeed the proximal cause of anomalous Pulfrich percepts, we presented identical stimuli in the two eyes and independently damped the onscreen amplitudes of the left and right eye motion trajectories. Again, anomalous Pulfrich percepts occur as expected directions. Then, we conducted an experiment with multiple levels of onscreen damping and measured psychometric functions. This experiment allowed us to estimate the relative neural damping caused by interocular differences in spatial frequency.

Finally, with a recently developed target-tracking paradigm for continuous psychophysics that uses hand movement as the measure of response, we demonstrate that the visuomotor system processes high spatial frequencies with more delay and longer temporal integration periods than low spatial frequencies. These results dovetail with those from the traditional forced-choice experiments. Previous studies have shown that delays in sensory processing are faithfully preserved in the movement of the hand (Burge & Cormack, 2020). (Similar findings have been reported for the smooth pursuit eye movements of the oculomotor system; Lee, Joshua, Medina, & Lisberger, 2016). We conclude that differences in temporal integration between the eyes can cause anomalous Pulfrich percepts.

## Results

We conducted four separate experiments to test the hypothesis that mismatched temporal integration periods can cause anomalous Pulfrich percepts. We used a within-subjects design. The first three experiments used a traditional forced choice paradigm. Observers binocularly viewed an oscillating Gabor stimulus and indicated the perceived orientation of its motion trajectory in depth. The fourth experiment was conducted using continuous target-tracking psychophysics (Bonnen, Burge, Yates, Pillow, & Cormack, 2015; Bonnen, Huk, & Cormack, 2017; Burge & Cormack, 2020; Knöll, Pillow, & Huk, 2018; Mulligan, Stevenson, & Cormack, 2013). Observers manually tracked a randomly moving Gabor stimulus with a cursor. The results of all experiments support the conclusion that different temporal integration periods can cause differential motion damping in the two eyes, and that this differential damping is the cause of anomalous Pulfrich percepts.

### Experiment 1: Neural damping with mismatched spatial frequencies in the two eyes

Experiment 1 was designed to establish the dependence of the anomalous Pulfrich effect on spatial frequency. On each trial, the observer was dichoptically presented an oscillating Gabor stimulus. The onscreen disparities specified a near-elliptical trajectory in depth that was aligned with the screen.

A different carrier spatial frequency was presented to each eye. Under our working hypothesis, the Gabor with the higher spatial frequency should be processed with more delay and (crucially) with a longer temporal integration period than the lower frequency Gabor in the other eye. The longer temporal integration period should cause damping of the effective motion in that eye. The damping, in turn, should cause the illusory orientation of the motion trajectory in depth. We predict that observers will report more ‘left-side-back’orientations when the left eye has the lower spatial frequency, and more ‘right-side-back’orientations when the right eye has the lower spatial frequency (Fig. 3AB).

**Figure 3.**
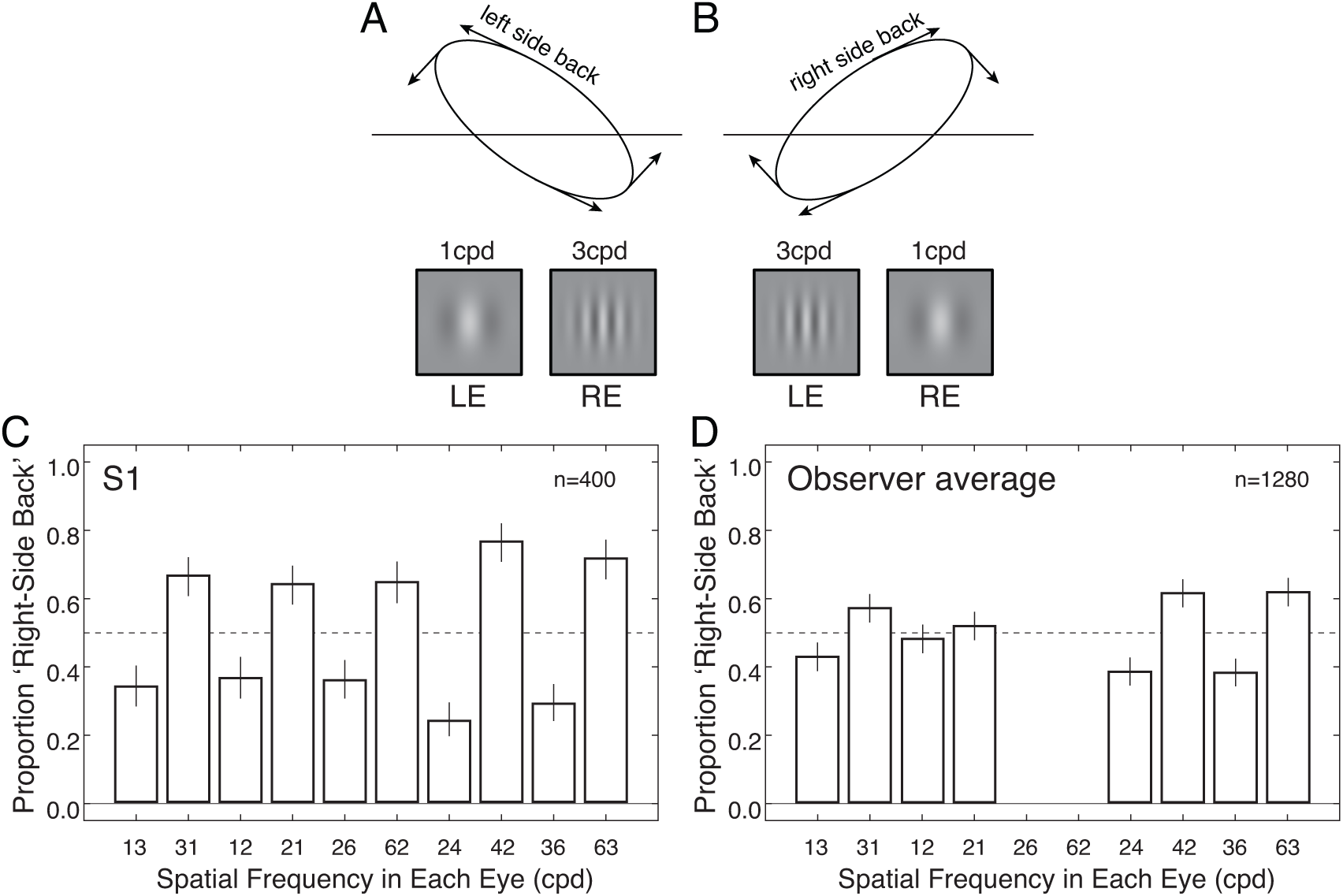
Experiment 1 stimuli, conditions, and results. **A** A low frequency Gabor in the left eye and a high frequency Gabor in the right eye predicts that the target will be perceived as moving along a trajectory oriented left-side-back in depth. **B** A high frequency Gabor in the left eye and a low frequency Gabor in the right eye predicts that the target will be perceived as moving along a trajectory oriented right-side-back in depth. Although fusion was imperfect, all observers reported percepts of motion in depth. **C** Mean-centered results from one observer (see Methods). A different spatial frequency was presented to each eye. In all cases, ‘right-side back’orientations were reported less often when the low frequency was in the left eye, and more often when the low frequency was in the right eye and. **D** Results combined across observers. Note that the 2cpd vs. 6cpd condition is absent from this plot. In the screening phase (see Methods), no observers other than observer S1 were able to fuse the stimuli in this condition, even when large onscreen disparities were present.

Observers were asked to report the apparent orientation of the motion trajectory in depth by indicating with a key press whether the principal axis of the trajectory appeared rotated left-side-back or right-side-back from the plane of the screen. Recall that the onscreen disparities specified that the motion trajectory was aligned with the screen. Absent eye-specific effects of spatial frequency, observers should not perceive left- and right-side-back orientations, such that each response is equally probable. If, on the other hand, if the two spatial frequencies are processed with different temporal integration periods, observers should report more ‘right-side-back’orientations when the right eye has the lower spatial frequency, and vice versa.

Observers reported more ‘right-side back’orientations of the perceived motion trajectory in depth when the right eye contained the lower frequency and more ‘left-side back’orientations when the left eye contained the lower frequency (Fig. 3CD; also see Fig. S1). In one observer, the effect appeared in all four conditions. For three observers, this effect was present in three out of the four conditions. This pattern of results is consistent with the experimental hypothesis. The null hypothesis is that spatial frequency has no effect on perceived orientation. Thus, if the null hypothesis was correct, observers should have reported right-side back orientations 50% of the time regardless of whether the left or right eye was presented the higher spatial frequency. We performed binomial tests on the group data to determine whether the null hypothesis could be rejected (see Methods). The tests rejected the null hypothesis for interocular spatial frequency combinations of 1cpd vs. 3cpd (*p*<0.01), 2cpd vs. 4cpd (*p*<0.001), and 3cpd vs. 6cpd (*p*<0.001), but not 1cpd vs. 2cpd (*p*=0.25). These results are largely consistent with the hypothesis that higher frequencies are processed by the visual system with longer temporal integration periods.

Conditions in which the left eye was stimulated with the lower spatial frequency (e.g. 1cpd to the left eye, 3cpd to the right eye; Fig. 3A) were interleaved with conditions in which the right eye was stimulated the lower spatial frequency (e.g. 3cpd to the left eye, 1cpd to the right eye; Fig. 3B). There are two benefits to this design. First, idiosyncratic block-specific response biases that may be present on a given block should be equally distributed amongst both conditions and have little effect on the final results. Second, because humans are poor at utrocular discrimination, it is difficult to determine which of the two eyes are being presented a given stimulus (Blake & Cormack, 1979; Schwarzkopf, Schindler, & Rees, 2010). Intermixing conditions ensures that, on any given trial, observers were unclear about which eye was being presented which stimulus. Hence, it would be quite difficult for observers to deliberately respond in a manner consistent with the experimental hypothesis.

Presenting substantially different images to the two eyes in a task that is supported in part by stereopsis raises concerns about poor binocular fusion. We screened eight observers for their ability to fuse and perform a stereo-motion task with mismatched stimuli in the two eyes (see Methods). Four of the eight screened observers were able to perform the task. That is, four of eight observers reported seeing depth and were able to acceptably fuse the target when large binocular disparities were presented onscreen.

### Experiment 2: Onscreen damping with matched spatial frequencies in the two eyes

In Experiment 1, our hypothesis is that the effective motion amplitude in one eye was damped because a higher spatial frequency was presented to that eye. Assuming the hypothesis is correct, Experiment 1 therefore manipulated motion amplitude indirectly. Experiment 2 was designed to provide direct evidence that damping the motion signal in one eye changes the perceived orientation of the motion trajectory in depth with respect to the screen. Experiment 2 is distinguished from Experiment 1 by two major design changes. First, the motion amplitude was damped onscreen rather than manipulated indirectly via interocular spatial frequency differences (Fig. 4AB). The resulting onscreen disparities specified a motion trajectory in depth with a principal axis that was misaligned with the screen. Unlike in Experiment 1, identical Gabors—with the same spatial frequencies used in Experiment 1—were presented to the two eyes. Matched Gabors alleviate the fusion difficulties associated with mismatched Gabors. More importantly, because the Gabors were matched, neural processing delays and temporal integration periods should be matched between the eyes.

**Figure 4.**
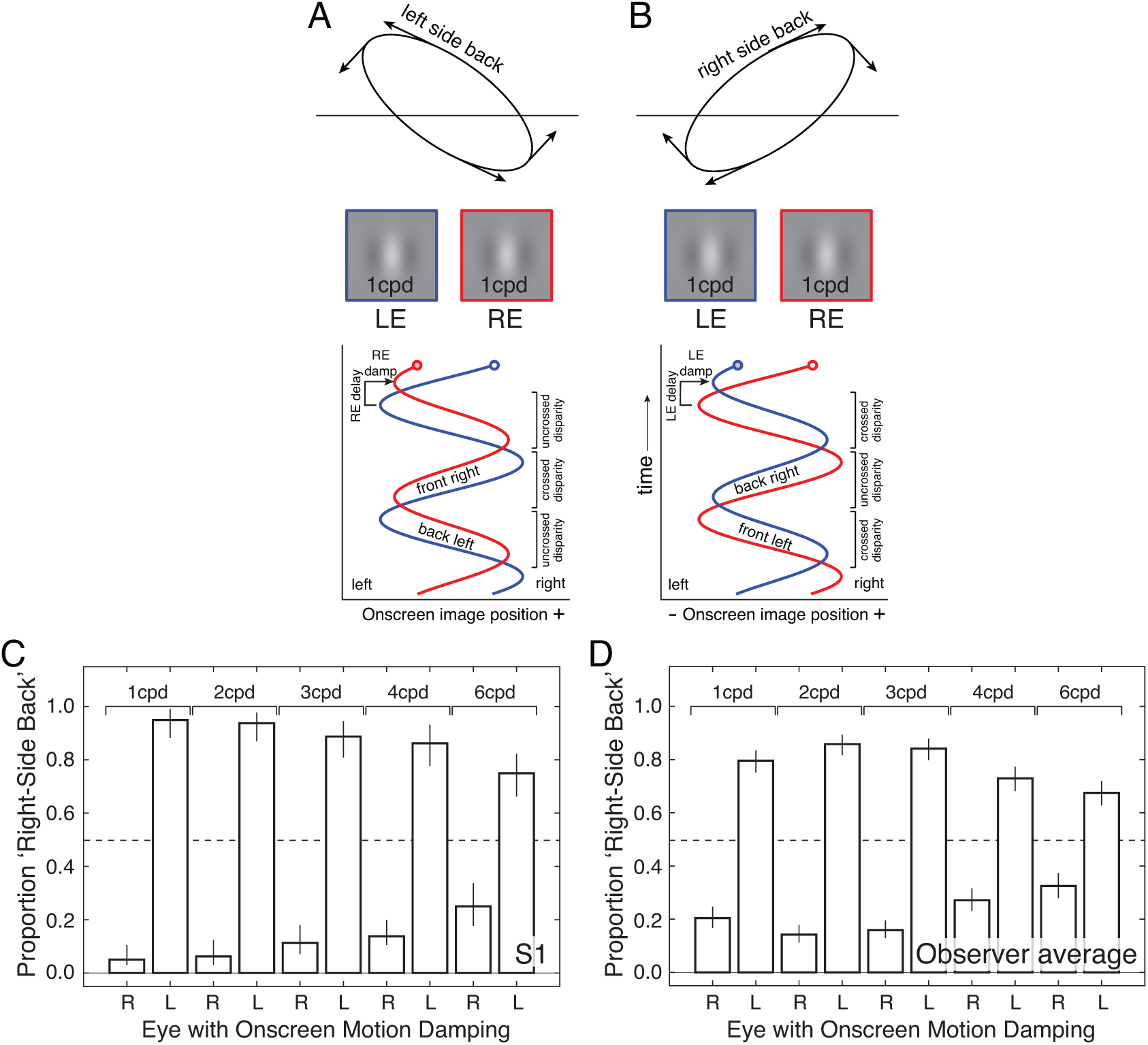
Experiment 2 stimuli, conditions, and results. **A** When the eyes are presented Gabors with the same spatial frequency, but the right-eye motion amplitude is damped (and delayed) onscreen, stereo-geometry specifies a near-elliptical motion trajectory that is oriented left-side back with respect to the screen. **B** When the eyes are presented Gabors with the same spatial frequency, but the left-eye motion amplitude is damped (and delayed) onscreen, stereo-geometry specifies a near-elliptical motion trajectory that is oriented right-side back with respect to the screen. **C** Mean-centered results from one observer. For all frequencies, responses are consistent with stereo-geometry-based predictions. Error bars indicate ±1 standard error. **D** Combined results across all observers.

The task performed by each observer was the same as in Experiment 1. Observers reported both whether the perceived motion trajectory appeared to be oriented ‘left-side back’or ‘right-side back’with respect to the screen. When the right-eye onscreen motion amplitude was damped relative to the left-eye onscreen motion amplitude, observers more often reported trajectories that were oriented left-side back. When the left-eye onscreen motion amplitude was damped relative to the right-eye motion amplitude, observers more often reported trajectories that were oriented right-side back (Fig. 4CD; also see Fig. S2). This data is similar to that collected in the first experiment. Recall that, under the working hypothesis, the mismatched Gabors in Experiment 1 should yield mismatched temporal integration periods between the eyes. The mismatched temporal integration periods, in turn, cause differential neural damping of the effective motion amplitude in the two eyes. The fact that the data exhibits similar patterns in the two experiments suggests that similar percepts result from differential onscreen damping, on one hand, and differential neural damping that results from mismatched Gabors in the two eyes, on the other.

### Experiment 3: Estimating the magnitude of neural damping

Experiment 3 was designed to measure the amount of neural damping that is induced by mismatched frequencies in the two eyes. The logic of the design is as follows: If anomalous Pulfrich percepts are due to neural damping differences in addition to neural delays, it should be possible to find the onscreen damping differences that eliminate the perceived orientation of the motion-in-depth trajectory. These onscreen damping differences should be equal in magnitude but opposite in sign of the neural damping differences that are induced by mismatched spatial frequencies in the two eyes.

We collected psychometric functions with onscreen damping difference as the independent variable in each condition, and measured the proportion of times that observers reported motion trajectories that were oriented ‘right side back’with respect to the screen. The resulting psychometric functions are shown in Fig. 5A (also see Fig. S3). The point of subjective equality (PSE) in each condition indicates the onscreen damping that is equal in magnitude and opposite in sign to the corresponding neural damping. When the right eye had the higher spatial-frequency stimulus, the left-eye onscreen motion had to be damped to eliminate anomalous Pulfrich percepts, and vice versa (Fig. 5BC; also see Fig. S4). This finding held across all tested frequency combinations. These results further support the hypothesis that stimulus-induced differences in temporal integration periods cause neural motion damping that can be neutralized by onscreen motion damping.

**Figure 5.**
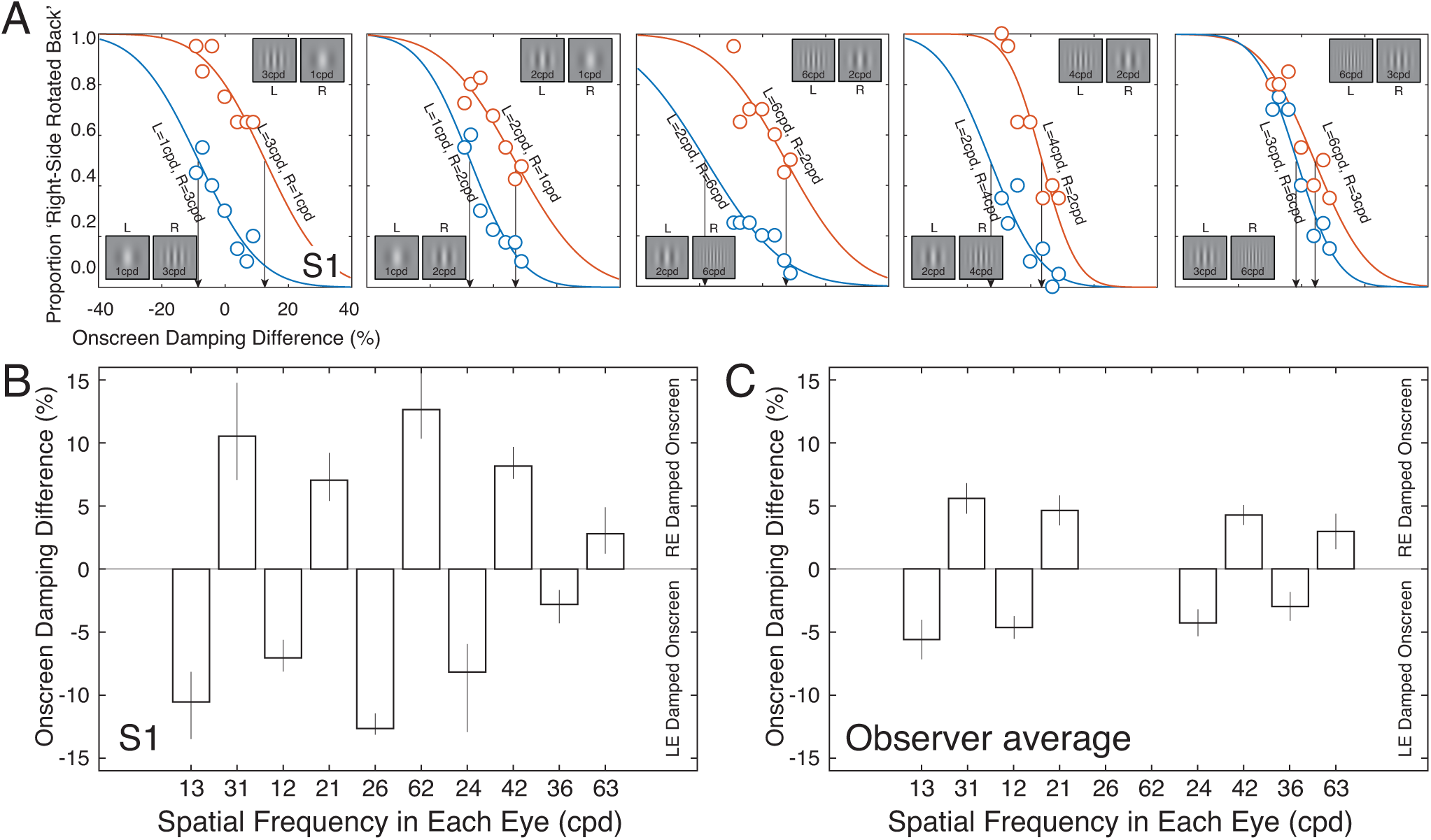
Experiment 3 stimuli, conditions, and results. **A** Psychometric functions from the first human observer for five different frequency pairs (i.e. 1cpd vs 3cpd, 1cpd vs 2cpd, 2cpd vs 6cpd, 2cpd vs 4cpd, and 3cpd vs 6cpd). When the left eye had the lower frequency stimulus, the psychometric functions were shifted consistently to the left (blue points and curves), indicating that the left-eye motion amplitude had to be damped onscreen to null the perceived orientation in depth. When the right eye had the lower frequency stimulus (red points and curves), the psychometric functions were shifted consistently to the right, indicating that the right-eye motion amplitude had to be damped onscreen to null the perceived orientations in depth. **B** Mean-centered points of subjective equality (PSEs; arrows in A) in each condition for one observer (see Methods). The PSEs are estimates of the amount of onscreen damping required to null the perceived orientations associated with different spatial frequencies in the two eyes. **C** Mean-centered PSEs averaged across all observers. Error bars represent bootstrapped ±1 standard errors on the points of subjective equality. Note that the 2cpd vs. 6cpd conditions are absent from this subplot. In the screening phase (see Methods), no observers other than observer S1 were able to perform the task in these conditions.

### Experiment 4: Continuous target-tracking psychophysics

The results from the first three experiments i) establish that mismatched spatial frequencies in the two eyes cause anomalous Pulfrich percepts, ii) demonstrate that damping the onscreen motion amplitude in one eye causes anomalous Pulfrich percepts with matched spatial frequencies in the two eyes, and iii) show that onscreen damping can eliminate spatial-frequency-induced anomalous Pulfrich percepts. Together, these results suggest that different temporal integration periods between the eyes are the root cause of spatial-frequency-induced anomalous Pulfrich percepts. The evidence for this conclusion, however, is indirect. To gain more direct evidence that mismatched frequencies induce interocular differences in temporal integration, Experiment 4 made use of an entirely different paradigm: continuous target-tracking psychophysics (Bonnen et al., 2015).

Using a mouse, observers manually tracked one of five Gabor targets at a time (Fig. 6A). The Gabor targets had carrier spatial frequencies of 1cpd, 2cpd, 3cpd, 4cpd, and 6cpd. These spatial frequencies were matched to those used in the previous experiments. For example, spatial frequencies of 1cpd and 3cpd were used in the target-tracking task because conditions in the previous experiments involved presenting a 1cpd Gabor to one eye and a 3cpd Gabor to the other. Throughout each run, the target underwent a horizontal random walk on the screen (Fig. 6B). The task was performed without difficulty. The cross-correlation between the target and response motions provides information about the temporal processing of the visuo-motor system. If the visuomotor system is linear, the cross-correlogram equals an estimate of the temporal impulse response function when the target velocities are white noise, which they are here by design.

**Figure 6.**
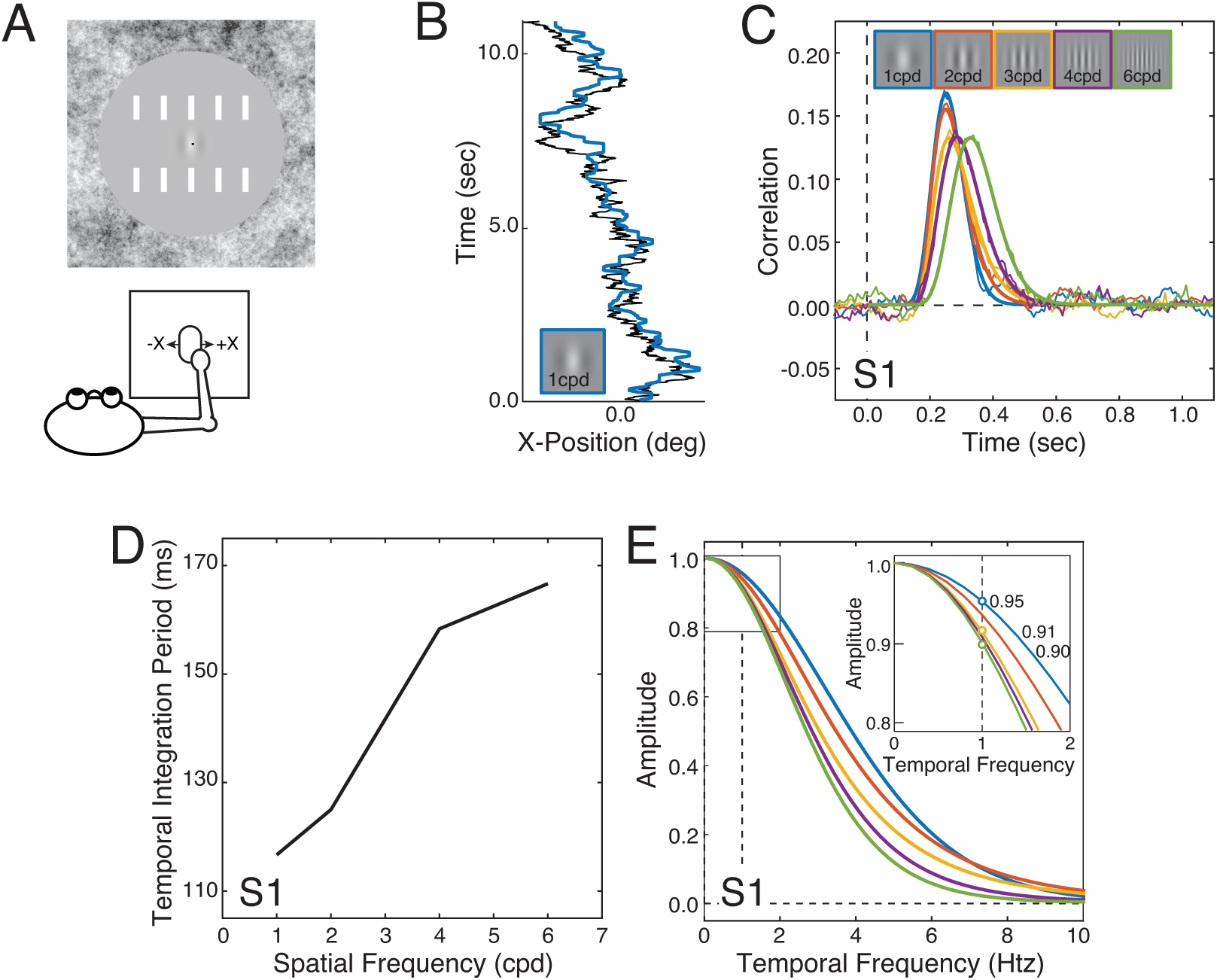
Effects of spatial frequency on target tracking performance for the first human observer. **A** On each trial, the observer tracked, with a mouse cursor (black dot at the center of the screen), a Gabor stimulus following a horizontal random walk across the center of the screen. **B** Example target-tracking performance on a single trial. The solid black trace indicates the horizontal random walk taken by the stimulus (1cpd Gabor). The blue trace indicates the position of the observer’s cursor. **C** Cross-correlograms in the target tracking task derived from target-tracking performance. The cross-correlograms change systematically as a function of spatial frequency (colors). **D** The temporal integration period (i.e. full-width at half-height) increases from approximately 115ms to 165ms as spatial frequency increases from 1cpd to 6cpd. **E** The amplitude spectra of the cross-correlograms provide an estimate of the amount of effective motion damping for each of many temporal frequencies. The inset shows the estimated amount of visuomotor motion damping for each spatial frequency at 1.0htz, the temporal frequency of the motion stimulus in the 2AFC experiments.

The cross-correlograms are broader in time as spatial frequency increases (Fig. 6CD). The amplitude spectra of the cross-correlograms indicate the proportion by which each spatial frequency is damped as a function of temporal frequency (Fig. 6E, see Methods). The inset shows at 1.0htz—the temporal frequency at which targets oscillated in the 2AFC forced-choice experiments (i.e. Exp. 1-3)—the motion amplitude of the visuomotor response in the tracking task decreases as spatial frequency increases. (The same is true for other temporal frequencies.) In other words, the amplitude of the visuomotor response is damped increasingly more as target spatial frequency increases.

To examine whether the visuomotor damping estimated in the target tracking task (Exp. 4, see Fig. 6E) can predict the effective sensory-perceptual motion damping estimated in the 2AFC forced-choice task (Exp. 3, see Fig. 5), we plotted the estimates of damping from the two experiments against each other. For the first human observer, the damping estimates are strongly correlated (r=0.90; p<0.04; Fig. 7A). The group average shows a similar trend (r=0.98; p=0.02; Fig. 7B). For all observers but one, the same qualitative pattern exists: sensory-perceptual- and tracking-based estimates of motion damping increase together. However, the slopes of the best-fitting lines vary substantially across observers (Fig. 7C). It will therefore be difficult, on an observer-by-observer basis, to predict the magnitude of the visuomotor motion damping in the tracking task from estimates of visual motion damping in the forced-choice task, or vice versa (see Discussion). Nevertheless, these results are consistent with the hypothesis that effective motion damping underlies anomalous Pulfrich percepts.

**Figure 7.**
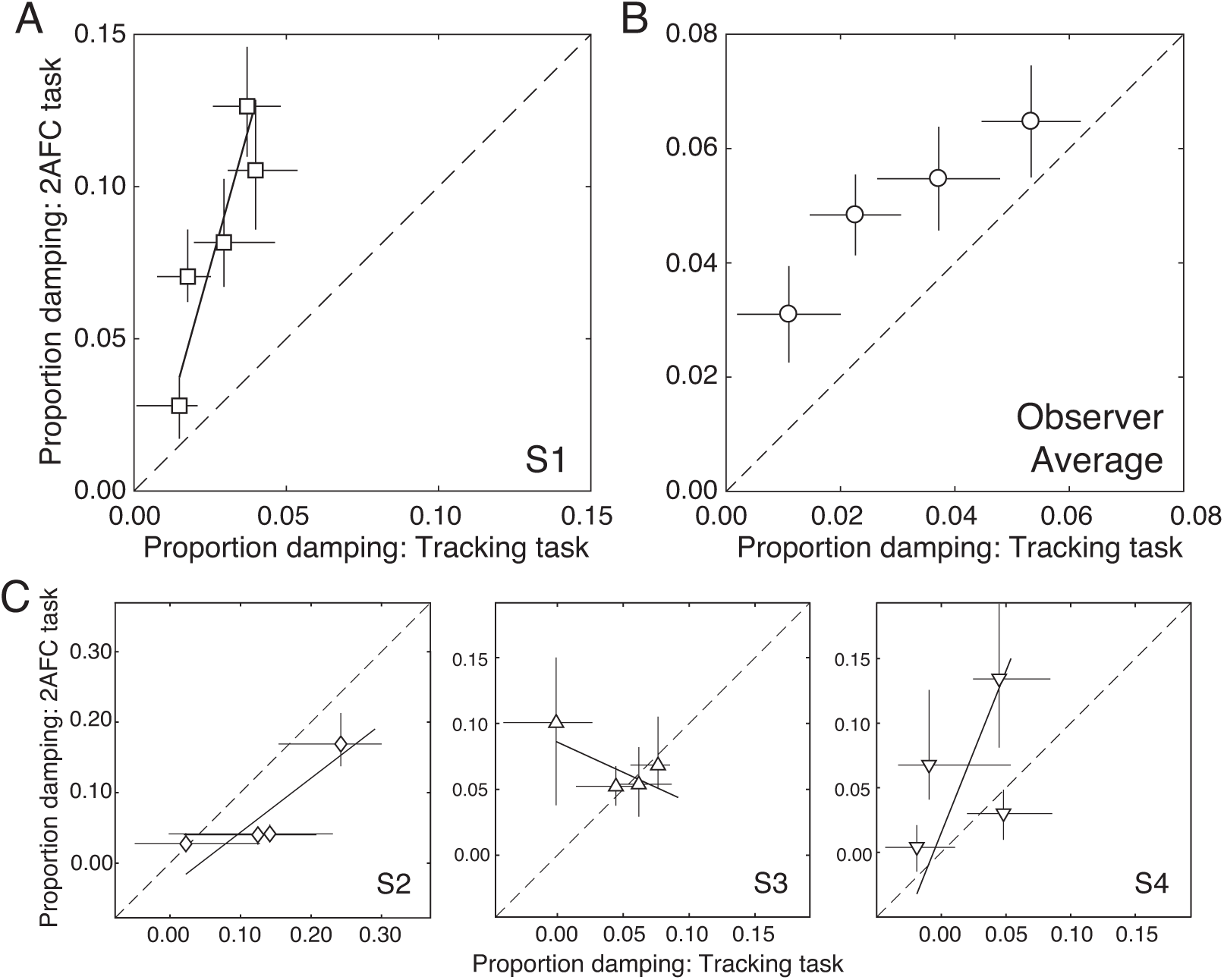
Comparison of 2AFC-based vs. target-tracking-based estimates of motion damping. **A** Results for the first observer. The best-fit line via weighted linear regression (solid), and the unity line (dashed) are also shown. Error bars on data points indicate 68% bootstrapped confidence intervals. **B** Average results across all observers (see Methods). **C** Results for each of the other three observers.

## Methods

### Participants

Four human observers participated in the experiment. Three observers were male and one observer was female. One was an author and the rest were naïve to the purposes of the experiment. All had normal or corrected to normal visual acuity (20/20), and normal stereoacuity as determined by the Titmus Stereo Test. The observers were aged 23, 26, 27, and 42 years old at the time of the measurements. All observers provided informed consent in accordance with the Declaration of Helsinki using a protocol approved by the Institutional Review Board at the University of Pennsylvania.

### Apparatus

Stimuli were presented on a custom four-mirror stereoscope. Left- and right-eye images were presented on two identical Vpixx VIEWPixx LED monitors. The monitors were 52.2×29.1cm, with a spatial resolution of 1920×1080 pixels, a refresh rate of 120Hz, and a maximum luminance of 105.9cd/m^2^. After light loss due to mirror reflections, the maximum luminance was 93.9cd/m^2^. The gamma function of each monitor was linearized using custom software routines. A single AMD FirePro D500 graphics card with 3GB GDDR5 VRAM controlled both monitors to ensure that the left and right eye images were presented simultaneously. To overcome bandwidth limitations of the monitor cables, custom firmware was written so that a single color channel drove each monitor; the red channel drove the left monitor and the green channel drove the right monitor. The single-channel drive to each monitor was then split to all three channels for gray scale presentation.

Observers viewed the monitors through a pair of mirror cubes positioned one inter-ocular distance apart. The mirror cubes had 2.5cm openings. Given the eye positions relative to the openings, the field of view through the mirror cubes was ∼15×15º. The outer mirrors were adjusted such that the vergence distance matched the 100cm distance of the monitors. This distance was confirmed both by a laser ruler measurement and by a visual comparison with a real target at 100cm. At this distance, each pixel subtended 1.09arcmin. Stimulus presentation was controlled via the Psychophysics Toolbox-3 (Brainard, 1997). Anti-aliasing enabled sub-pixel resolution permitting accurate presentations of disparities as small as 15-20arcsec. Heads were stabilized with a chin and forehead rest.

### Forced-choice psychophysics target motion

For the forced-choice psychophysics experiments, we simulated the classic pendulum Pulfrich stimulus on the display. For each trial, the left- and right-eye on-screen bar positions in degrees of visual angle were given by

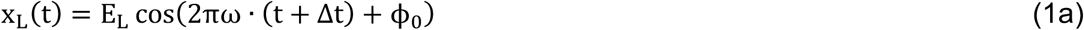

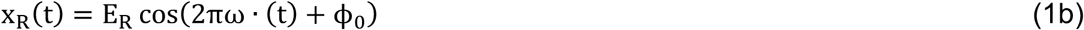

where E_L_ and E_R_ are the left- and right-eye motion amplitudes in degrees of visual angle, Δ*t* is the on-screen delay between the left- and right-eye target images, ω is the temporal frequency of the target movement, *ϕ*_0_ is the starting phase, and *t* is time in seconds.

The undamped motion amplitude was 1.5º of visual angle (3.0º total change in visual angle in each direction). The maximum onscreen motion damping in one eye (20%) corresponded to 80% (1.2º of visual angle) of the undamped amplitude in the other. The range of particular damping values was adjusted to the sensitivity of each observer. The on-screen interocular delays were set at ±25ms. The temporal frequency was 1 cycle per second. The starting phase *ϕ*_0_ was randomly chosen on each trial to equal either 0 or *π*, which forced the stimuli to start either to the left or to the right of center.

When the onscreen interocular difference in motion amplitude equals zero and the onscreen interocular delay is zero, the target moves in the frontoparallel plane at the distance of the screen; the onscreen disparities are zero throughout the trial. If the interocular difference in motion amplitude is non-zero and/or if the interocular delay is non-zero spatial binocular disparities result, and the disparity-specified target follows a motion-in-depth trajectory outside the plane of the monitor. Differences in motion amplitude cause a disparity-specified misalignment in depth of the motion trajectory. Non-zero delays cause a disparity-specified elliptical trajectory of motion in depth. Negative delay values indicate the left-eye on-screen image is delayed relative to the right; positive delay values indicate the left eye on-screen image is advanced relative to the right.

The on-screen binocular disparity for a given interocular delay and damping as a function of time is given by

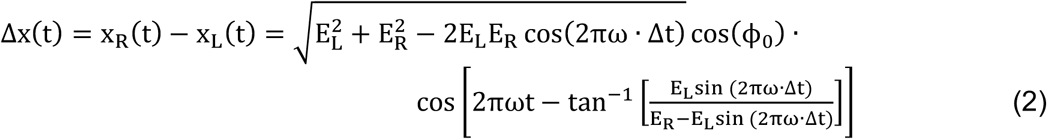

where negative disparities are crossed (i.e. nearer than the screen) and positive disparities are uncrossed (i.e. farther than the screen). The disparity takes on its maximum magnitude when the perceived stimulus is directly in front of the observer and the lateral movement is at its maximum speed. The maximum disparity in visual angle is given by 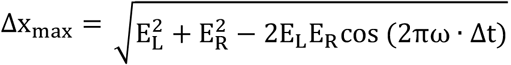 and it occurs when 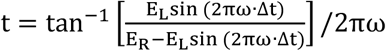. Note that we did not temporally manipulate when left- and right-eye images were presented on-screen; both eyes’images were presented coincidently on each monitor refresh. Rather, we calculated the disparity 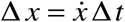 given the target velocity and the desired on-screen delay on each time step, and appropriately shifted the spatial positions of the left- and right-eye images.

Two sets of five vertically-oriented picket-fence bars (0.25×1.00º) flanked the region of the screen traversed by the target stimulus. The picket fences were specified by disparity to be at the screen distance. A 1/f noise texture, also defined by disparity to be at the screen distance, covered the periphery of the display. Both the picket fences and the 1/f noise texture served as stereoscopic references to the screen distance and helped to anchor vergence.

Before the target appeared on each trial, a small dot appeared 1.5º to the left of center or 1.5º right of center at the location of imminent target appearance. Observers were instructed to fixate the dot and then, after the target appeared, fixate and follow the target throughout the trial. In pilot experiments, we found that if observers did not follow the target with their eyes, the highest spatial frequency Gabor occasionally appeared to vanish during the trial at and near when it hit top speed (i.e. 6cpd). Observers reported whether the perceived motion trajectory was oriented left-side back or right-side back from frontoparallel. All experiments used a one-interval, two-alternative forced choice procedure.

### Forced-choice psychophysics stimuli

The same Gabor targets were presented in the forced-choice psychophysical experiments as in the tracking experiments. A vertically-oriented Gabor is given by the product of a sinewave carrier and a Gaussian envelope

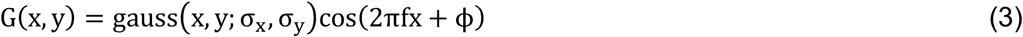

where σ_x_ and σ_y_ are the standard deviation in X and Y of the Gaussian envelope, f is the frequency of the carrier, and ϕis the phase. Five Gabor targets with different carrier frequencies were used: 1cpd, 2cpd, 3cpd, 4cpd, and 6cpd. All had the spatial size because all had same Gaussian envelope (σ_x_ = 0.39 and σ_y_ = 0.32). The octave bandwidths thus equaled 1.5, 0.7, 0.46, 0.35, 0.23 and the orientation bandwidths equaled 60º, 32º, 22º, 16º, and 11º, respectively. The phase of the carrier frequency was equal to 0.0 for all Gabor stimuli (i.e. all Gabors were in cosine phase).

Experiment 1 presented Gabors with different spatial frequencies in the two eyes. Data was collected in blocks with an intermixed design. For example, blocks containing conditions in which the left and right eyes were respectively presented 1cpd and 3cpd Gabors were intermixed with conditions in which the left and right eyes were presented 3cpd and 1cpd Gabors. In each condition, we used two values of interocular delay. The increased neural temporal integration period associated with high spatial frequencies served to dampen the effective motion amplitude in one eye relative to the other. Human observers have poor utrocular discrimination; humans have significant difficulty determining which eye is being presented a given stimulus (Blake & Cormack, 1979; Schwarzkopf et al., 2010). Intermixing conditions ensured that, on any given trial, observers—even non-naïve observers—were unclear about which eye was being presented which stimulus. Thus, observers—even non-naïve observers—would be unable to determine, on a given trial, which response was consistent with the experimental hypothesis.

Experiment 2 presented Gabors with the same spatial frequency in the two eyes, and used two interocular delays and two damping values. We chose damping values that made the orientation of the near-elliptical trajectory in depth (‘left side back’vs. ‘right side back’) easy for the observers to identify.

Experiment 3 was designed to measure observer sensitivity to interocular differences in motion amplitude (i.e. damping). Experiment 3 thus measured full psychometric functions in each condition using the method of constant stimuli. Seven different levels of damping were collected for each function. The psychometric functions were fit with a cumulative Gaussian via maximum likelihood methods. The 50% point on the psychometric function—the point of subjective equality (PSE)—indicates the onscreen motion damping needed to null the relative motion damping due to spatial frequency differences. Observers ran 140 trials per condition (i.e. 140 trials per psychometric function) in counter-balanced blocks of 70 trials each.

### Mean-centering of effects

Data from Experiments 1-3 were mean-centered for pairs of matched conditions. Matched conditions were those involving the same spatial frequencies (e.g. 1cpd vs. 3cpd and 3cpd vs. 1cpd). The proportion of ‘right-side back’responses or effective damping was mean-centered across matched conditions according to the following equation:

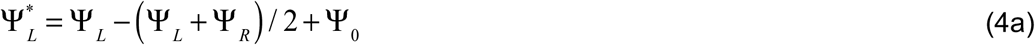

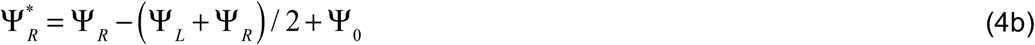

For Exp. 1, Ψ represents the proportion of ‘right-side back’responses, where Ψ _*L*_ and Ψ _*R*_ respectively correspond to conditions where the left eye, or the right eye, were presented the lower spatial frequency. For Exp. 2, Ψ also represents the proportion of ‘right-side back’responses, and Ψ _*L*_ and Ψ _*R*_ respectively correspond to conditions where the onscreen motion was damped in the left eye, or in the right eye. For Exp. 3, Ψ represents the psychophysical estimate of onscreen motion damping, 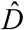, that is required to null the neural damping, and Ψ _*L*_ and Ψ _*R*_ correspond to the condition in which onscreen motion was damped in the left-eye, or in the right eye, respectively. In Experiments 1 and 2, Ψ_0_ has a value of 0.5. In Experiment 3, Ψ_0_ has a value of 0%.

### Binomial test for significance

Under our working hypothesis, ‘right-side back’responses should be reported more often when the effective motion-amplitude in the left eye is smaller than that in the right eye. Similarly, when the effective motion-amplitude in the right eye is smaller than that in the left eye, ‘right-side back’responses should be reported less often. The null hypothesis predicts that there will be no difference in the proportion of ‘right-side back’responses across two matched conditions (e.g. 1cpd vs. 3cpd and 3cpd vs. 1cpd). To determine whether the proportions of ‘right-side back’responses differed significantly from those predicted by the null hypothesis, we used a binominal test. Under the null hypothesis, the probability, *p*, of the observed response proportions is given by

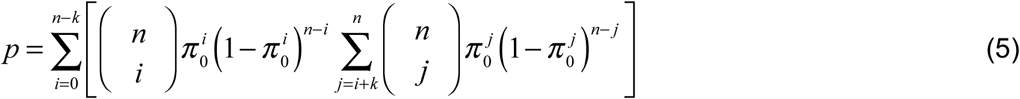

where *n* is the number of trials in a given condition, *π* _*o*_ is the probability of the observer responding ‘right-side back’in each of the two matched conditions under the null hypothesis (i.e., 0.5), and *k* is the difference in the number of ‘right-side back’responses between two matched conditions.

### Reliability-weighted averaging of estimated motion damping

PSEs estimates in Experiment 3 were averaged across observers using reliability-weighted averaging:

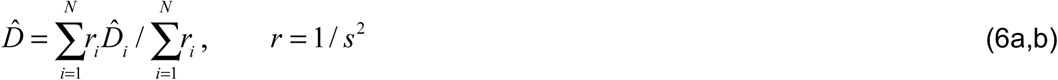

where 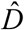 is the estimate of motion damping for a given condition, averaged across all observers. *N* is the number of observers and *s* is the standard error of motion damping estimates (as determined by 68% bootstrapped confidence intervals). Reliability-weighted averaging takes into consideration differences in the reliability of damping estimates across observers. These differences in reliability arise because some observers are more sensitive to onscreen motion damping than others. It is well-known from signal detection theory that greater sensitivity in a task is associated with more reliable estimates of the point of subjective equality (here, estimates of motion damping).

### Estimated relationship between forced-choice- and target-tracking-based motion damping estimates

The relationship between 2AFC-based and target-tracking-based estimates of motion damping was fit with a line via weighted linear regression. Since estimates of motion damping from both tasks have associated uncertainty, simple linear regression is not appropriate. This is because simple linear regression assumes that one of the variables is independent, and thus has no associated uncertainty. We fit the parameters of the best-fit line with maximum likelihood methods using numerical optimization. The cost function was

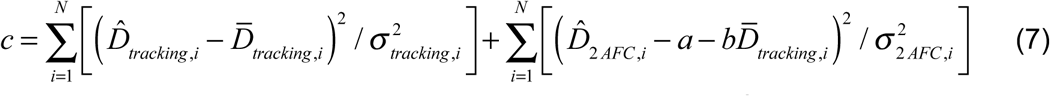

where *N* is the total number of conditions for an observer, 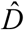 is the experiment-derived estimate of motion damping for a given condition, 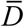 is a free parameter indicating the expected amount of motion damping for a given condition, *σ* is the standard error of the motion damping estimate for a given condition (as determined by 68% bootstrapped confidence intervals), *a* is the y-intercept of the best fit line, and *b* is the slope of the best fit line.

### Observer screening

Before inclusion in the main experiments, observers were screened for their ability to perform the task when the spatial frequencies in the two eyes differed by a factor of three. During this screening phase, the onscreen motion amplitude differed in the two eyes by a large amount of up to 20%. These onscreen amplitude differences caused the stereo-specified motion trajectory to be misaligned with the screen. If an observer was unable to correctly report the direction of the stereo-specified misalignment at least 80% of the time, no further data was collected from that observer. Four out of eight screened observers were excluded from the study on this basis. The excluded observers all reported difficulty fusing and difficulty seeing any stereo-specified depth at all. The pilot data is consistent with these reports.

### Target-tracking procedure

Tracking data was collected from each observer in blocks of individual runs. Each run was initiated with a mouse click, which caused the target and a small dark mouse cursor to appear in the center of the screen. After a stationary period of 500ms, the target began a one-dimensional horizontal random walk (i.e. Brownian motion) for eleven seconds. The task was to track the target as accurately as possible with a small dark mouse cursor. Blocks contained intermixed runs from each of the four conditions.

### Target-tracking psychophysics: Onscreen stimuli

Data was collected in five conditions, each of which was distinguished by a different target Gabor stimulus. Each Gabor target had one of five different carrier frequencies: 1cpd, 2cpd, 3cpd, 4cpd, and 6cpd. All Gabor targets shared the same Gaussian envelope (*σ*_*x*_=0.39º & *σ*_*y*_=0.32º), and subtended approximately 2.0ºx2.0º of visual angle (i.e. five sigma). Hence, in the five conditions, the octave bandwidths equaled 1.5, 0.7, 0.46, 0.35, and 0.23 and the orientation bandwidths equaled 60º, 32º, 22º, 16º, and 11º, respectively. Data was collected in five intermixed blocks of twenty runs each for a total 20 runs per condition.

### Target-tracking psychophysics: Target motion

For the tracking experiments, the target stimulus performed a random walk on a gray background subtending 10.0×7.5º of visual angle, and was surrounded by a static field of 1/f noise. The region of the screen traversed by the target was flanked by two horizontal sets of thirteen vertically-oriented picket fence bars (Fig. 6A).

The x-positions of the target on each time step *t* + 1 were generated as follows

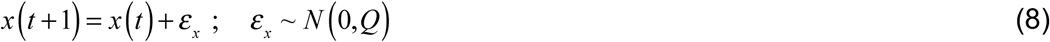

where *ε* _*x*_ is a random sample of Gaussian noise and *Q* is the drift variance. The random sample determines the change in target position between the current and the next time step. The drift variance determines the expected magnitude of the position change on each time step, and hence the overall variance of the random walk. The variance of the walk positions across multiple walks *σ* ^2^ (*t*) = *Qt* is equal to the product of the drift variance and the number of elapsed time steps. The value of the drift variance in our task (0.8mm per time step) was chosen to be as large as possible such that each walk would traverse as much ground as possible while maintaining the expectation that less than one walk out of 500 (i.e. less than one per human observer throughout the experiment) would escape the horizontal extent of the gray background area (176×131mm) before the 11 second trial completed.

The effective on-screen positions of the images are obtained by convolving the on-screen target images with the temporal impulse response function

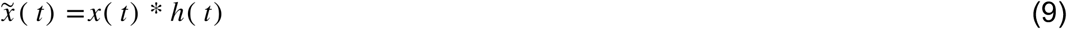

where *h*(*t*) is a temporal impulse response function corresponding to a specific frequency. Convolving the target velocities with the impulse response function gives the velocities of the effective target images. Integrating these velocities across time gives the effective target positions.

To determine the impulse response function relating the target and response, we computed the zero-mean normalized cross-correlations between the target and response velocities

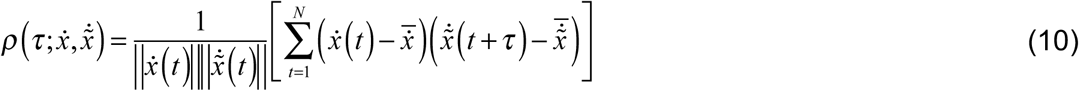

where *τ* is the lag, 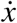 and 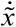 are the target and response velocities. Assuming a linear system, when the input time series (i.e. the target velocities) is white, as it is here by design, the cross-correlation with the response gives the impulse response function of the system.

To compute the normalized cross-correlations, we did not include the first second of each eleven second tracking run so that observers reached steady state tracking performance. The mean cross-correlation functions shown in the figures were obtained by first computing the normalized cross-correlation in each run (Eqn. 10), and then averaging these cross-correlograms across runs in each condition.

### Gamma distribution fits to mean cross-correlograms

To summarize the mean cross-correlograms, we fit a Gamma distribution function using maximum likelihood methods. The form of the fitted function was given by

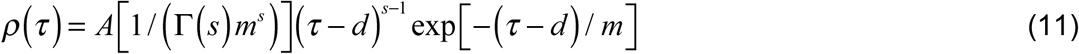

where *A* is the amplitude, and *m, s*, and *d* are the parameters determining the shape and scale of the fit. The mode (i.e. peak) of the function is given by *ms*. We use the mode as our measure of delay. The full-width at half-height can be used as a measure of the temporal integration period, and can be computed via numeric methods. The damping associated with a given fitted function is given by the value of the normalized amplitude spectrum at the temporal frequency of the stimulus, which in the current experiments is one cycle per second.

## Discussion

In this manuscript, we presented evidence that anomalous Pulfrich percepts—illusory motion trajectories in depth misaligned with the true direction of motion—are caused by interocular differences in temporal integration periods in the two eyes. This specific perceptual effect, and the reasons it occurs, have more general implications.

The integration of multiple complementary streams of incoming information with different temporal dynamics is fundamental to the performance of biological systems. In most cases, sensory-perceptual systems successfully solve this temporal binding problem, and compute accurate estimates of environmental properties. In some cases, the visual system fails to compensate for temporally mismatched signals, and inaccurate estimates result. Such cases are instructive. They can help reveal fundamental properties about the temporal nature of sensory signals, and make plain the striking perceptual consequences of insufficient compensatory mechanisms.

In this discussion section, we contextualize the anomalous Pulfrich effect with reference to other areas of vision research, consider how visual and visuomotor measures of performance are related, and discuss potential future directions.

### Analogy to the Geometric effect in surface orientation perception

Horizontal minification (or magnification) of the image in one eye causes the misperception of surface orientation. This phenomenon is known as the Geometric effect (Banks & Backus, 1998; Ogle, 1950). The Geometric effect occurs because the horizontal minification in one eye distorts the patterns of binocular disparity such that they specify a surface slant that is different from the actual surface slant. For example, when a frontoparallel surface is viewed with a horizontal minifier in front of the right eye, the surface is perceived to be slanted left-side back. If the left-eye image is minified, the same surface is perceived to be slanted right-side back (Fig. 8).

**Figure 8.**
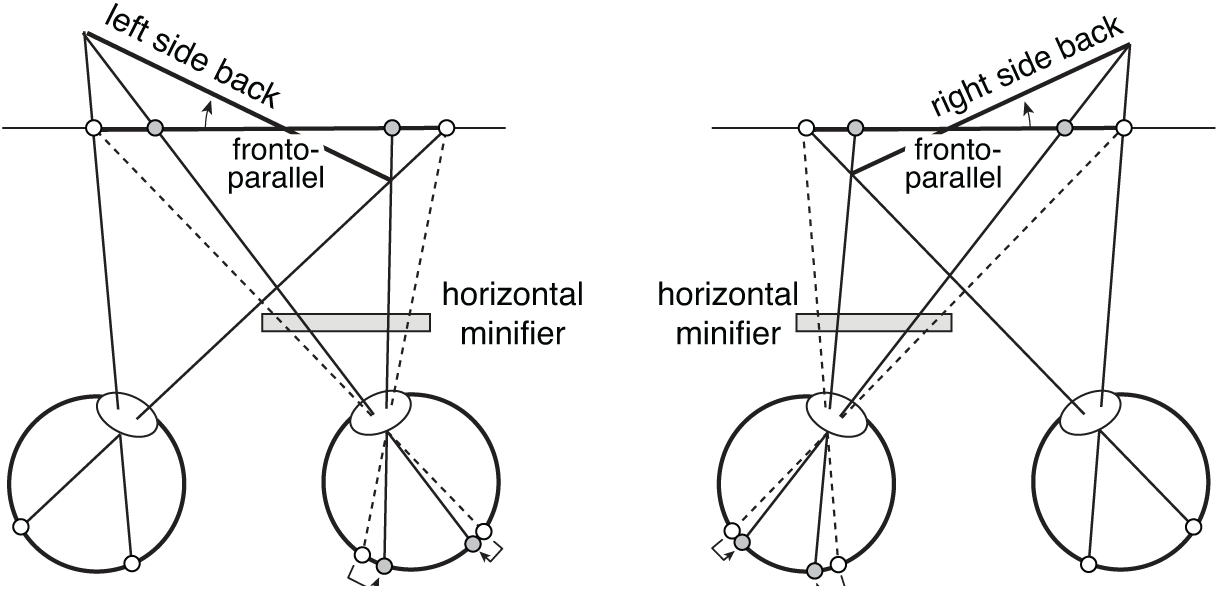
The Geometric effect in stereo-slant perception. Horizontal minification (or magnification) distorts the pattern of binocular disparities such that the disparity-specified orientation of the surface appears rotated in depth. If the horizontal minifier is in front of the right eye, a frontal surface straight-ahead is perceived left side back. If the horizontal minifier is in front of the left eye, a frontal surface straight-ahead is perceived right side back. The same principles account for both the Geometric effect and anomalous Pulfrich percepts.

The principles behind the Geometric effect mirror the principles behind the anomalous Pulfrich effect. An obvious analogy can be drawn between right- or left-eye motion damping and right- or left-eye horizontal minification. Anomalous Pulfrich percepts are caused by motion that is differentially damped between the two eyes. Indeed, if the effective image motion is damped but not delayed in one eye relative to the other, the disparity-specified motion trajectory lies in the plane of the slanted surface specified by disparities caused by the Geometric effect.

### Preservation of sensory processing dynamics in motor movements

The current manuscript reports a series of results that strongly suggest that different spatial frequencies are processed with different temporal integration periods, and that these differences underlie anomalous Pulfrich percepts. Linking the target-tracking results to sensory-perceptual processing requires an assumption. The assumption is that changes in the ability of an observer to track a target across different target stimuli reflect changes in the sensory-perceptual processing of the stimuli as opposed to changes in the motor response. Multiple studies have shown this assumption holds in various situations. Motor variation in smooth-pursuit eye movements is due overwhelmingly to sensory errors (Osborne, Lisberger, & Bialek, 2005). Changes in the width of the cross-correlogram associating target and hand movements during target-tracking are linked to the sensitivity of visual target location discrimination (Bonnen et al., 2015). Delays in visual processing match delays in the motor response of both the eye (Lee et al., 2016), and the hand (Burge & Cormack, 2020; Lee et al., 2016). However, it appears from the present experiments that differences in the visual temporal integration period are not always faithfully preserved in the motor response of the hand.

Experiments 1-3 used traditional forced-choice psychophysical techniques to establish the anomalous Pulfrich phenomenon and quantify the effective motion damping that is caused by differences in temporal processing induced by different spatial frequencies. Experiment 4 used continuous target-tracking psychophysics to collect more direct evidence that different spatial frequencies are indeed associated with different temporal integration periods. The average estimates of motion damping across human observers from the target-tracking task very nearly matched those from the forced-choice task (see Fig. 7B). But there was significant inter-observer variability regarding how the two sets of estimates were related (see Fig. 7A,C). In two of four observers, the forced-choice-based estimates were systematically larger than the tracking-based estimates. In one observer, the reverse was true. And in the remaining observer, the estimates were nearly matched, except for an apparent outlier.

The finding that forced-choice- and target-tracking-based estimates of damping are correlated but do not exactly agree for individual observers warrants further study. Our analysis assumes that the motor component of the visuomotor response can be accurately modeled with convolution, a linear open-loop computation. It is likely that there are benefits to modeling visuomotor performance in the target-tracking task as a closed-loop system, given that visual feedback is integral to good performance in many visuomotor tasks. It is also possible that convolution does not accurately capture how the motor system translates visual input into a motor response. If so, other (possibly nonlinear) operations will be required to accurately model the motor contribution to performance. These, and related, issues are under active investigation.

### Computational challenges of mismatched temporal processing

The visual system must constantly deal with the problem of staggered information arrival. We have focused on the perceptual consequences of temporal processing differences associated with mismatched spatial frequency content in the two eyes. Interocular differences in spatial frequency content commonly occur in natural viewing. During binocular viewing of surfaces that are slanted about a vertical axis, for example, the spatial frequencies tend to be higher in one eye than the other. These differences, while extremely common, tend to be relatively small. For a surface at a distance of 30cm and a slant of 72º, the corresponding frequencies in the two eyes will differ by approximately a factor of two (i.e. horizontal size ratios of 0.5 or 2.0, depending on whether the surface is slanted left- or right-side back). For more distant and less slanted surfaces, which are more common in natural viewing (Adams et al., 2016; Backus, Banks, van Ee, & Crowell, 1999; Burge, McCann, & Geisler, 2016; Kim & Burge, 2018; 2020; Yang & Purves, 2003), the ratio tends to be substantially smaller. However, typical natural images have broadband 1/f spectra, and frequencies above the contrast detection threshold typically vary by a factor of ten or more. Thus, the temporal binding problem may be a more acute computational challenge within each eye’s image than between the images in the two eyes. In spite of this challenge, the visual system usually generates (largely) accurate estimates of environmental properties.

Measuring the temporal processing constraints of the nervous system, and developing normative theory for how different streams of information should be integrated to achieve accurate perceptual estimates, will help advance our understanding of how the spatial-frequency binding problem is resolved by biological systems (Burge et al., 2019). Incorporating these solutions into image-computable ideal observers for sensory-perceptual tasks with natural stimuli is a potentially fruitful future direction for neuroscience and vision research (Burge, 2020; Burge & Geisler, 2011; 2012; 2014; 2015; Chin & Burge, 2020).

## Conclusion

The problem of binding temporally damped and temporally staggered information is not a niche problem. It is not at all specific to the combination of information from different spatial frequency channels, as we have focused on in this paper. The visual system must resolve temporal differences between luminance and chromatic signals, high and low luminance signals, and high and low contrast signals. More generally, the different senses—visual, auditory, vestibular, proprioceptive, tactile—transmit signals possessing substantially different temporal properties. These signals must also be combined to form accurate, temporally coherent percepts. Future work will investigate how sensory-perceptual systems solve the temporal binding problem within and across senses.

## Supplement

**Figure S1.**
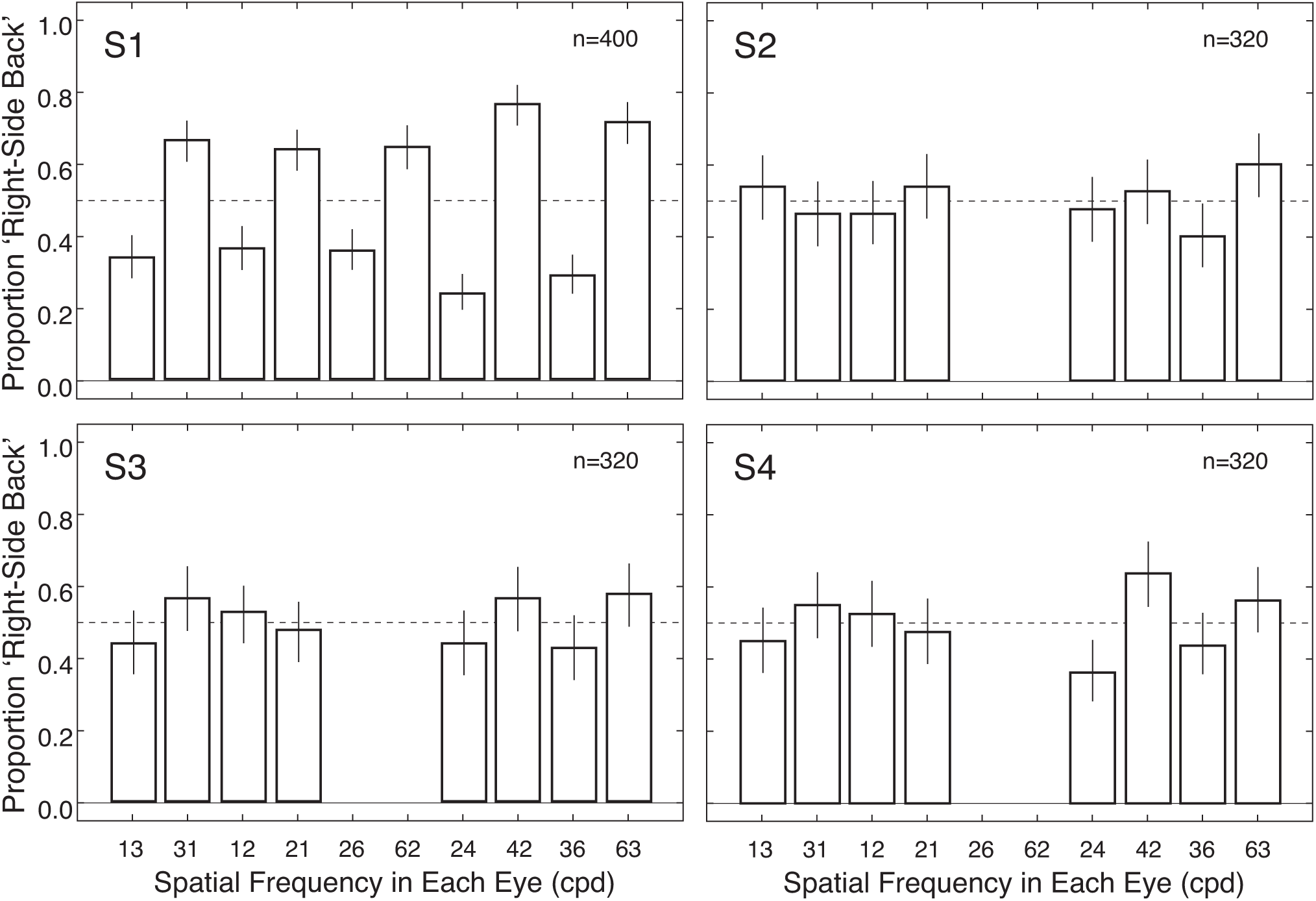
Experiment 1 results for all observers and conditions.

**Figure S2.**
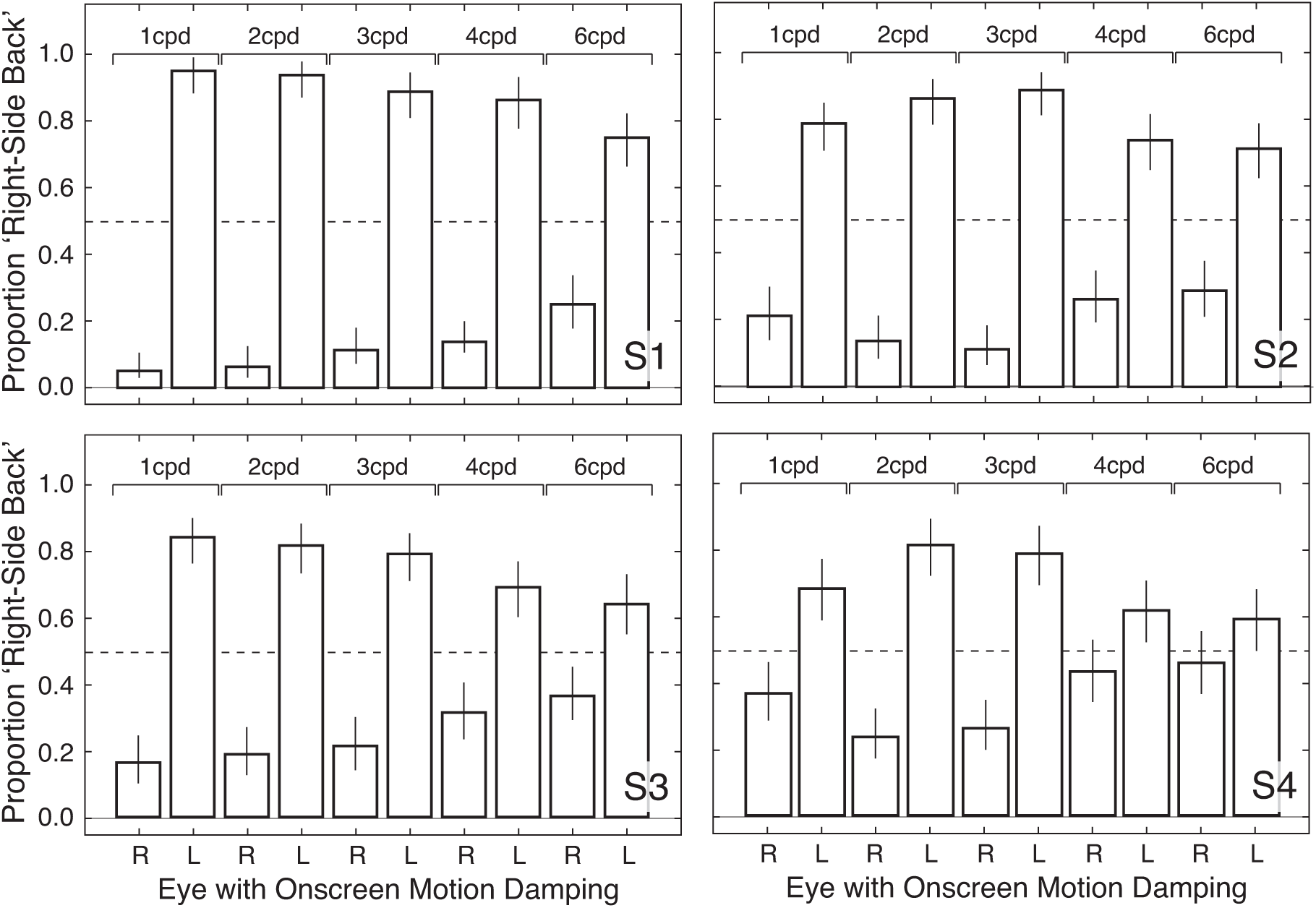
Experiment 2 results for all observers and conditions.

**Figure S3.**
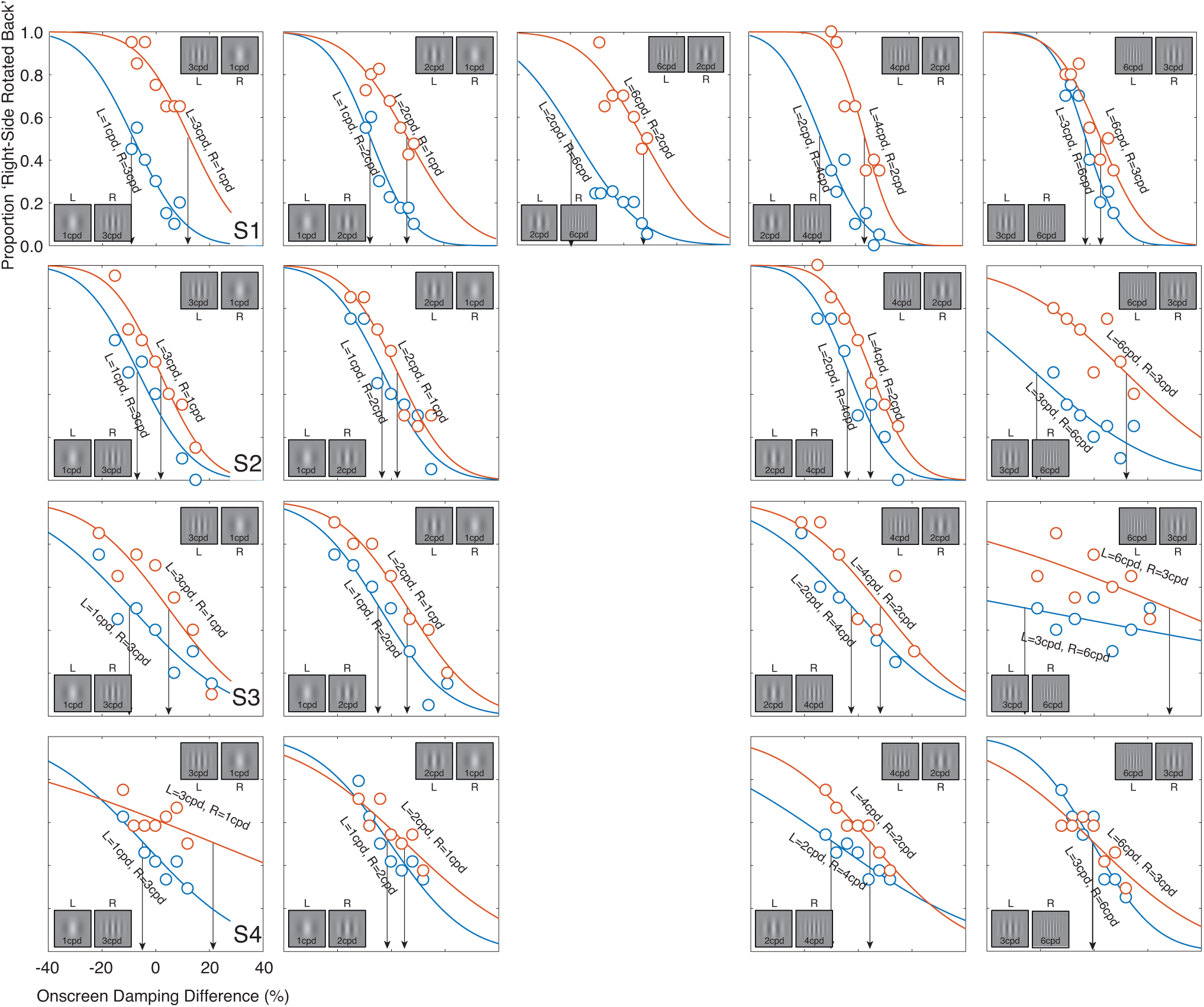
Experiment 3 stimuli, conditions, and psychometric functions for all observers and conditions. The data from all four observers follow the same qualitative pattern.

**Figure S4.**
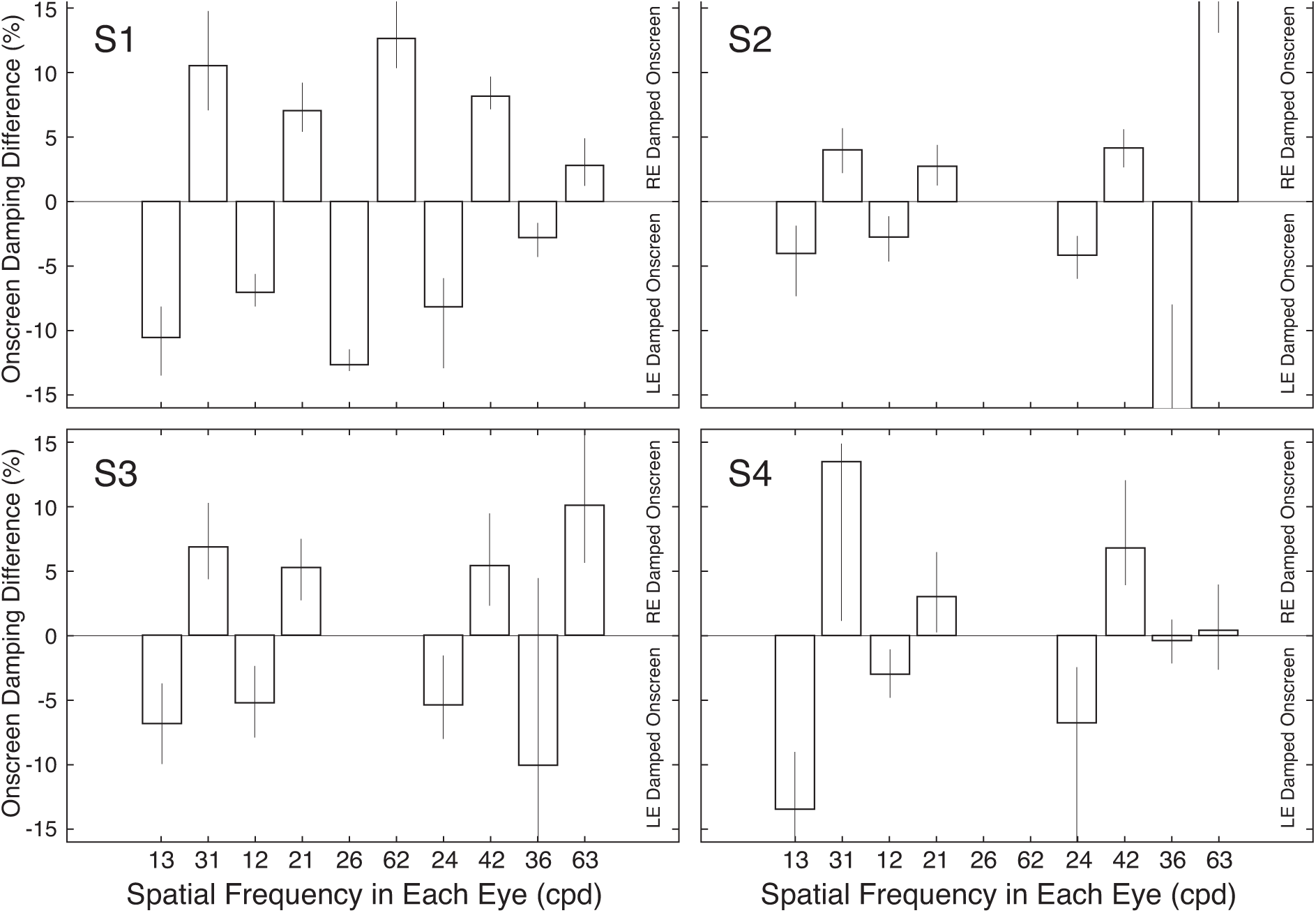
Experiment 3 points of subjective equality (PSEs) for all observers and conditions. The data from all four observers follow the same qualitative pattern. Note that for observer S2, the PSEs for the rightmost two conditions are larger than the scale of the plot.

**Figure S5.**
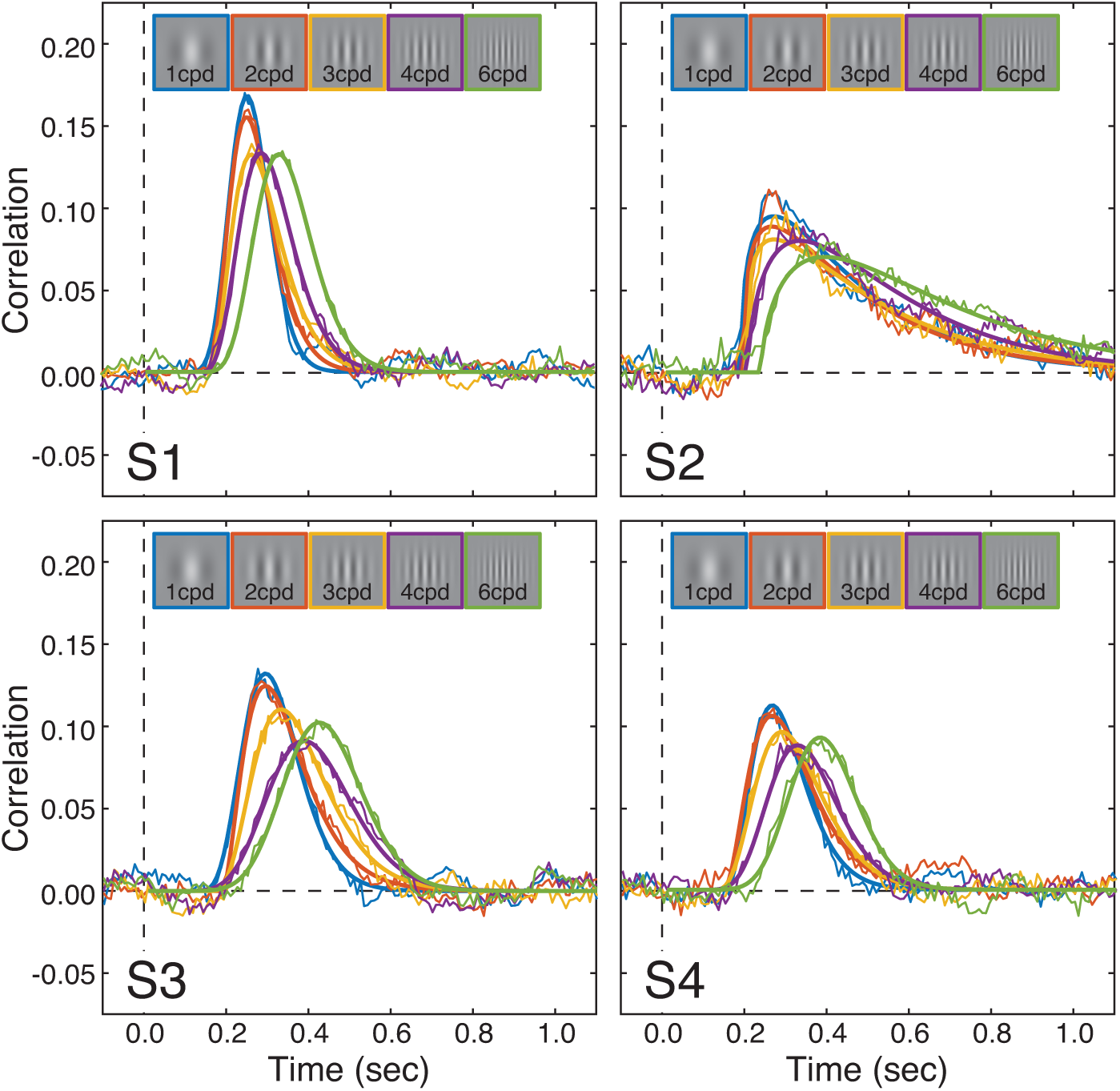
Experiment 4 stimuli and cross-correlograms for all observers and conditions.

